# Mass Spectrometry-Guided Library preparation of Peptides as Metabolic enzyme Natural Inhibitors for the management of Obesity and Diabetes

**DOI:** 10.1101/2024.12.23.630156

**Authors:** Manish Singh Sansi, Daraksha Iram, Sudarshan Kumar, Suman kapila, Kamal Gandhi, Sunita Meena

## Abstract

Goat whey protein (GWP) is recognized as a valuable source of bioactive peptides with significant health-promoting properties. In this study, GWP was enzymatically hydrolyzed using a combination of gastrointestinal enzymes— pepsin, trypsin, and chymotrypsin—to generate peptides. These peptides were identified through high-resolution liquid chromatography-mass spectrometry (HR-LC/MS), resulting in library of 2,883 peptides with lengths ranging from 6 to 44 amino acids. Among them, 40 peptides were predicted to exhibit high bioactivity scores (0.90–1) based on PeptideRanker analysis, with 28 of these being classified as nontoxic. Molecular docking simulations were employed to investigate the interactions of these peptides with pancreatic lipase and α-amylase to screening of inhibitors, these two enzymes critical in lipid and carbohydrate metabolism. Several peptides demonstrated strong binding affinities, suggesting their potential as enzyme inhibitors. Notably, peptides WPGIMR and WQDGSWQF showed the highest binding affinity for pancreatic lipase, while AAPFIWL and WQDGSWQF exhibited significant interactions with α-amylase. These results shed light on the molecular mechanisms underlying the inhibitory activities of whey protein-derived peptides. They highlight their potential applications as functional food ingredients or natural therapeutic agents for managing metabolic disorders such as obesity and diabetes, advancing the understanding of whey protein hydrolysates in modulating key metabolic enzymes.

## 1.0 Introduction

Milk proteins are significant sources of bioactive peptides, which can promote health benefits beyond their nutritional value (Korhonen & Pihlanto, 2006). These peptides, encrypted within protein sequences, require enzymatic hydrolysis, typically with enzymes like subtilisin or trypsin, to release their bioactive forms characterized by hydrophobic or positively charged C-terminal amino acids (Hernández-Ledesma et al., 2011; Power et al., 2013). Regarding the protein substrates, most of the studies dealing with the production of bioactive peptides from milk proteins employ cow milk as a raw material (Nagpal et al., 2011). Goat milk, with a higher proportion of β-casein and superior digestibility compared to cow milk, has shown potential for producing bioactive peptides with antihypertensive, antioxidant, and cholesterol-lowering properties (Ahmed et al., 2015; Park, 2009; Muro Urista et al., 2011).

The increasing prevalence of metabolic disorders such as obesity and type 2 diabetes has become a pressing global health issue. These conditions are often linked to the excessive intake of calorie-dense foods, which can disrupt normal lipid and carbohydrate metabolism, resulting in adverse health effects (Klein et al., 2022).

Enzymes like pancreatic lipase and α-amylase play critical roles in breaking down dietary fats and carbohydrates, making them important targets for therapeutic strategies to address these disorders (Aguiar et al., 2024; Kaur et al., 2021).

Pancreatic lipase is a key enzyme in the digestion of triglycerides, converting them into absorbable components. Inhibiting its activity can reduce triglyceride absorption and limit caloric intake, a primary contributor to obesity (Patil et al., 2017; Bello et al., 2017). Medications such as orlistat, which inhibit pancreatic lipase, have demonstrated their effectiveness in promoting weight loss and improving blood sugar regulation. However, their use is often accompanied by side effects, including hyperoxaluria due to disrupted fat absorption, which underscores the need for safer, naturally derived alternatives (Buysschaert et al., 2016; Patil et al., 2017).

Similarly, α-amylase is responsible for initiating the digestion of starch by hydrolyzing internal α-1,4-glucosidic bonds, producing maltotriose, maltose, glucose, and dextrin. Rapid starch digestion can lead to sudden increases in blood glucose and insulin levels, which are significant risk factors for type 2 diabetes (Ramasubbu et al., 2003; Butterworth et al., 2011; Labes et al., 2008). Slowing carbohydrate digestion through α-amylase inhibition can help stabilize postprandial glucose levels and reduce the risk of insulin resistance and related complications (Dhital et al., 2017; Kato et al., 2017; Jayaraj et al., 2013). Recently, attention has shifted toward natural inhibitors of pancreatic lipase and α-amylase due to their potential to provide safer and more sustainable therapeutic options. Protein hydrolysates and peptide fractions from various natural sources have been shown to positively influence lipid and carbohydrate metabolism, offering anti-obesity benefits (Birari & Bhutani, 2007). These findings emphasize the importance of exploring natural bioactive compounds to develop innovative treatments for metabolic disorders. This research focuses on further investigating such natural alternatives to create effective management strategies for obesity and diabetes. Whey protein, a byproduct of cheese production, is a rich source of peptides with bioactive potential (Dinika et al., 2020; Verma et al., 2023). These peptides, released through enzymatic hydrolysis. The therapeutic potential of whey protein-derived peptides requires advanced proteomic tools to identify, characterize, and evaluate their functionality.

Recent advances in mass spectrometry and bioinformatics approaches have enabled the precise identification and functional prediction of bioactive peptides (Fabre et al., 2021). Additionally, molecular docking simulations provide insights into the binding interactions of these peptides with target enzymes, elucidating their mechanisms of action. In this study, gastrointestinal enzymes (pepsin, trypsin and chymotrypsin) for proteolytic enzymatic hydrolysis of goat milk whey protein were hydrolysed to release a diverse array of whey protein peptides (WPP). Further, the proteomic characterization and molecular mechanism of WPP as inhibitors of pancreatic lipase and α-amylase.

A combination of HR-LC/MS-based peptide library preparation and identification. Bioinformatics tools, such as PeptideRanker and ToxinPred, was employed to predict bioactivity. Molecular docking simulations were conducted to assess the binding affinity and interactions of selected peptides with pancreatic lipase and α-amylase inhibitors. This work aims to uncover the therapeutic potential of whey protein-derived peptides in modulating lipid and carbohydrate metabolism, presenting an innovative approach to address the growing challenge of metabolic disorders. By elucidating the molecular interactions of these peptides with key enzymes, this study contributes to the development of functional foods and nutraceuticals for improving metabolic health.

## 2.0. Methods and material

WP-PTC hydrolysate was purified and sequence of peptides were identified using high-resolution liquid chromatography-tandem mass spectrometry (HR-LC/MS). Bioinformatics analysis, using tools like the Trans-Proteomic pipeline (TPP), potential peptides were identified. Then, bioactivity of peptides was predicted by peptide ranker tool, followed by prediction of physiological properties of peptides (molecular weight, length of peptides). Thereafter, selected bioactive peptides targeted to α-amylase and pancreatic lipase active site to analyse the interaction of peptides by molecular docking (Autodockvina).

### 2.1. Milk Collection

Goat milk was procured from Livestock Research Center at ICAR-National Dairy Research Institute, Karnal, India. These animals were raised under a consistent feeding and breeding regimen as part of a larger herd at the institute. The milk samples were collected and pooled before the experiment. The aim of pooling the samples was to minimize animal-to-animal variations.

### 2.2. Partial purification of casein and whey proteins

To isolate the casein and whey proteins raw goat milk was skimmed using a centrifuge at 4255×g for 20 min at 10 □C. Afterward, an acid precipitation was performed on the skimmed goat milk using 1N HCl at pH 4.6 to separate the whey and caseins. Separation involved centrifugation at 4255 ×g for 10 min at 4 □C, which was repeated thrice to ensure maximum removal of casein particles. The separated caseins proteins (CP) were dissolved in double-distilled water using 0.1 M NaOH which was converted to Na-caseinate. Afterward, they were lyophilized with a lyophilizer and stored at −20 □C for future experiments. To prevent the proteolytic degradation of milk whey, 0.01% phenyl methyl sulfonate was added. To isolate the whey protein (WP), the salting-out method was used, followed by dialysis using 3kDa cut off membrane.

### 2.3. Acetone precipitation

Dialyzed WP were then additionally precipitated by adding 3-fold chilled acetone and incubated at −20 □C for 14–16 h. Protein precipitation was followed centrifugation at 13,000 x g at 4 □C for 10 min, after that supernatant was discarded. The pellet was reconstituted with 1x PBS at pH 6.8. After that, the vacuum-sealed concentrator was used to dry the partially purified WP, which was then kept at −20 □C for upcoming tests.

### 2.4. Protein profiling by SDS-PAGE

Before conducting the SDS-PAGE, the concentrations of partially purified CP and WP were determined by the Lowry’s method (Lowry et al., 1951). The molecular weight of the proteins was determined using SDS-PAGE, and the protein profile was confirmed. The separating gel, following Laemmli (1970) approach, was composed of 15%, and the stacking gel was made up of 4% polyacrylamide. Sample loaded in 5-10µg per well. After electrophoresis, the gels were stained with a solution of 0.005% Coomassie brilliant blue R-250 in a mixture of glacial acetic acid, methanol, and water in a ratio of 1:5:4. for 30 min. Following staining, the gels were fixed with 10% trichloroacetic acid (TCA) for further analysis. Gels were destained, and after that, gel were observed by image scanner system (E-Gel Imager).

### 2.5. In-solution digestion of whey protein by pepsin, trypsin and chymotrypsin combination

For in-solution digestion, whey protein (1mg/mL) were taken then precipitated using 3fold acetone or trichloroacetic acid (TCA). Furthermore, 45mM DTT (dithiothreitol) was dissolved in 50 mM NH4HCO3 (ammonium bicarbonate) to reduce disulfide bonds in the protein samples for 20 min. This was followed by the alkylation of cysteine residues using 10 mM IAA (iodoacetamide) in 50 mM NH4HCO3 (ammonium bicarbonate) for 15 min. Subsequently, digestion was carried out using pepsin (1:100) at pH 2.0. Pepsin was inactivated by raising the pH to 7.5-7.8 using 0.1 mM NaOH. Then, trypsin and chymotrypsin were added (1:100) and the mixture was incubated at 37 □C for three hours. The reaction was subsequently halted with 10% TFA (trifluoroacetic acid), and the peptides were further dried using vacuum drying. The samples were then desalted using a zip tip and subjected to peptide identification by HR-LC/MS Orbitrap (Ali et al., 2017).

### 2.6. Electrospray ionization tandem mass spectrometry HR-LC/MS orbitrap analysis

The lyophilized peptides were reconstituted in 0.1% formic acid in LC-MS-grade water and subjected to nano-LC (Thermo EASY-nLC; Thermo Fischer Scientific) followed by identification in captive ion source i.e. Q Exactive Plus-orbitrap mass spectrometer (MS) with high mass accuracy and sensitivity. The peptides were enriched in nano-trap column and eluted from nano-analytical column. The peptide elution was carried out using a linear gradient of 5–45% acetonitrile at 300 µL/min flow rate in a total run time of 90 min keeping the solvent system as follows: solvent A, 100% water in 0.1% formic acid; and solvent B, 100% acetonitrile in 0.1% formic acid. Positive ions were generated by electrospray, and the orbitrap was operated in data-dependent acquisition mode to automatically switch between MS and MS/MS acquisition. Precursor ion orbitrap survey scan was acquired with a range of 350–2,000 *m*/*z* with resolution R =70,000 @ m/z 200. Q1 sequentially selects six most intense precursor ions for fragmentation using collision-induced dissociation for MS-MS analysis with a fixed cycle time of 3 sec along with 2 min of release for exclusion filter. Data acquisition was conducted by software, version 24.8; Thermo Fischer Scientific.

### 2.7. Data processing and analysis

Vendor-created .raw file format was analyzed with TPP search engine which is based on comet algorithm, used for high confidence identification of the *de novo* peptides library. Resultant .raw files were used for the identification of proteins using the profiled inhouse combined goat database (UniprotKb proteome database) along with usual contaminant proteins for spectra examination. The parameter for MS/MS ion search contains digested sites with two missed cleavages allowed, flexible modification on amino acids-methionine for oxidation, N-terminal acetylation, Gln-pyro Glu, and Glu-pyro Glu, while static modification of cysteine as carboxy amidomethylation or propionamide. The peptide precursor ion tolerance was 20 ppm, with MS/MS tolerance of 0.01 Da. The “ion score cutoff” was set to 30, thereby eliminating the lowest quality matches and minimum peptide length as five amino acid residues. To increase the confidence and remove the false-positive identification, a 0.01% false discovery rate (FDR) was used at equal peptide and protein levels.

### 2.8. In silico prediction of length, bioactivity and molecular weight of identified peptides

*In silico* prediction of peptides with specific properties, such as molecular weight and length, involves computational analysis. The bioactivity of peptides is predicted using a peptide ranker. Furthermore, the physicochemical properties of peptide sequences, molecular mass, and including toxicity were determined through using the ToxinPred tool (Mooney et al., 2012.

### 2.9. Peptide binding interaction with pancreatic lipase and α-amylase

After predicting the bioactivity of peptides, 28 peptides were chosen for molecular interaction analysis with pancreatic lipase and alpha-amylase. Docking simulations were utilized to gain a deeper understanding of the interaction between the most effective bioactive peptides and the binding sites of pancreatic lipase (PDB id: 1LPB) and α-amylase (1JFH). PyRx (AutoDockVina) was employed to conduct the molecular docking investigations. Discovery Studio 2022 software was used to evaluate docking orientations and interactions with binding pocket residues.

## 3.0. Result and discussion

The graphical workflow provides a comprehensive overview of the process for profiling and identifying anti-obesity and antidiabetic peptides derived from whey protein hydrolysate peptides. The procedure begins with the collection of goat milk. After the milk is collected, initial sample processing is conducted to prepare the whey fraction for further processing. The next step involves the purification of the whey protein. This is accomplished through a combination of methods: salting out, acetone precipitation, and dialysis. Salting out uses high salt concentrations to precipitate proteins, which helps isolate the whey proteins from other soluble milk components. Acetone precipitation further purifies the proteins by selectively removing contaminants and concentrating the whey proteins. Dialysis is then employed to remove small molecular contaminants, such as salts or solvents, by using a semi-permeable membrane that allows only the passage of small molecules, leaving behind larger proteins. Once the whey protein is purified, it is subjected to enzymatic digestion using digestive endopeptidases, specifically pepsin, trypsin, and chymotrypsin. These enzymes break down the whey protein into smaller peptides by cleaving specific peptide bonds, producing a mixture of peptides with potential bioactive properties. These peptides are then purified using a Zip Tip C18 column, a solid-phase extraction method. Following the purification, the peptides are analyzed using high-resolution liquid chromatography-mass spectrometry (HR-LC/MS) for the generation of peptides library. This advanced analytical technique separates and identifies the peptides based on their mass-to-charge ratio and provides detailed information on their molecular structure and sequence. The peptide profiling is further refined through the use of an in-house proteome database of goat. The peptide sequences are compared against this database using the TPP (Trans-Proteomic Pipeline), a bioinformatics toolset that helps identify and quantify peptides and proteins by matching experimental data with theoretical sequences. This step ensures the accurate identification of peptides and enhances the confidence in the findings. Finally, the peptides library was subjected to bioinformatics analysis, where various computational tools was used to predict the bioactivity of the identified peptides, specifically focusing on their potential anti-obesity and antidiabetic effects. This process, as depicted in the graphical workflow (Fig 3.0), highlights the integrated approach used in profiling anti-obesity and antidiabetic peptides and emphasizes the importance of both experimental techniques and bioinformatics in identifying potential bioactive peptides.

**Figure 3.0.**
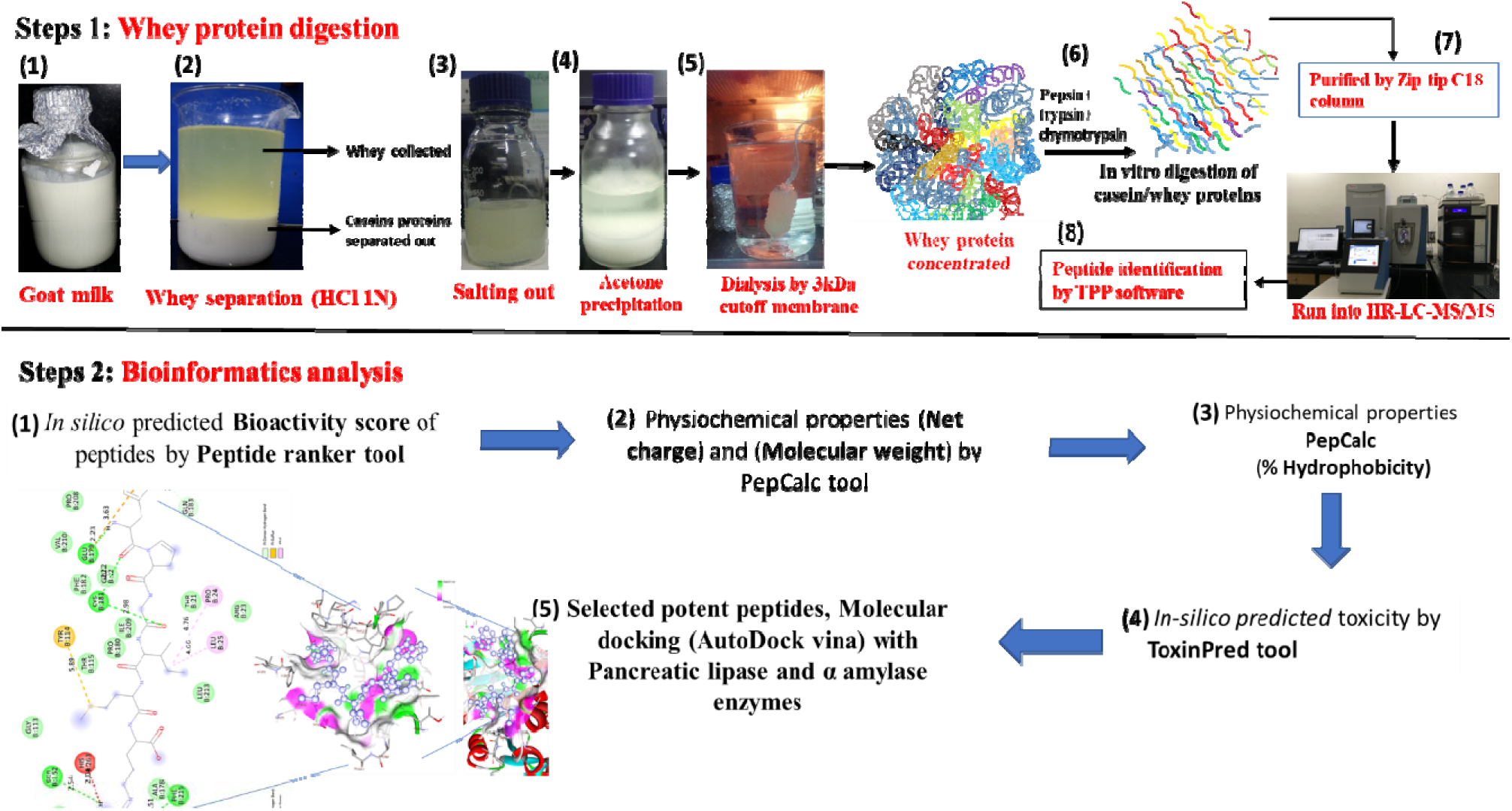
Workflow of profiling and identification of bioactive peptides from WP-PTC hydrolysate.

### 3.1. Nano-HPLC chromatogram and total ions spectra of HR-LC/MS of whey protein hydrolysate

Whey protein hydrolysate peptides was selected for identifying peptide sequences. HR-LC/MS was employed to determine the amino acid sequences and generated library of peptides, with the results depicted in Fig. 3.1 a & b. Subsequently, peptides were identified using an in-house combined database on the Trans-Proteomic-Pipeline (TPP) software. Utilizing the database to confirm amino acid sequences, a library of 2,883 peptides were identified in the whey protein hydrolysate peptides (Fig. 3.2. a, Supplementary-I). Moreover, it’s noteworthy that all peptides range from 6 to 44 amino acids in length, with the majority falling within the range of 8 to 13 residues (Fig. 3.2. a).

**Figure 3.1.**
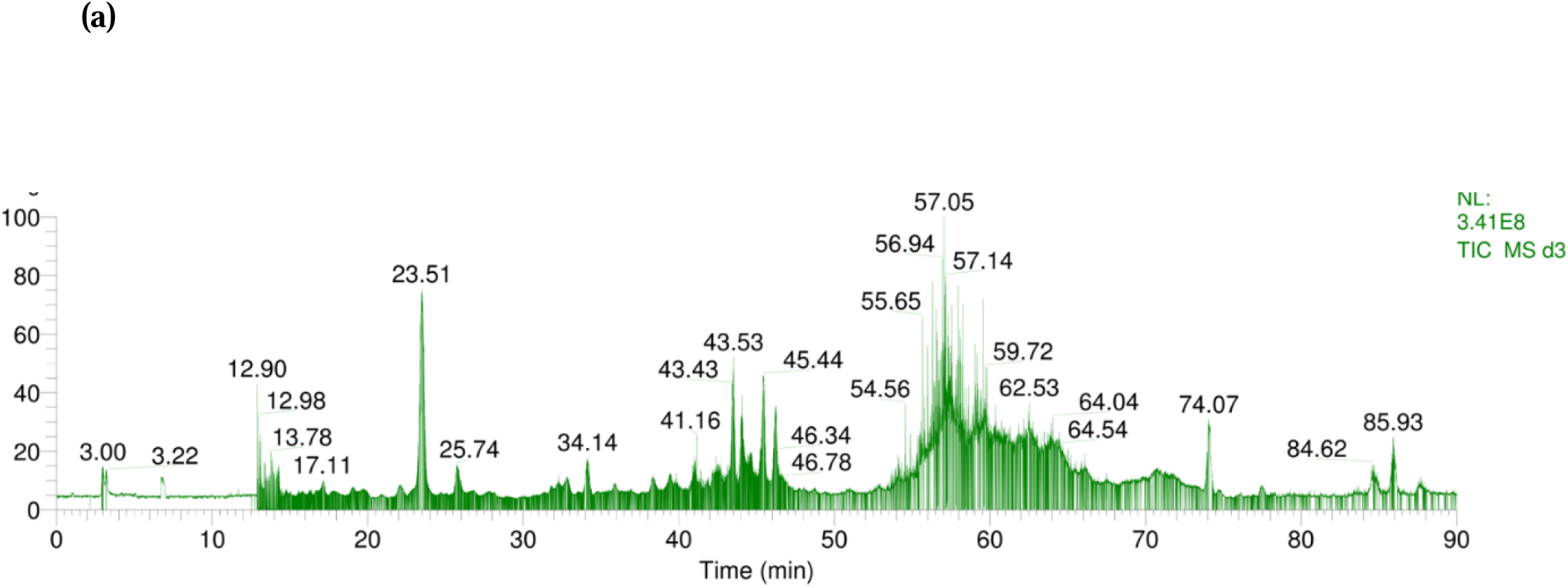
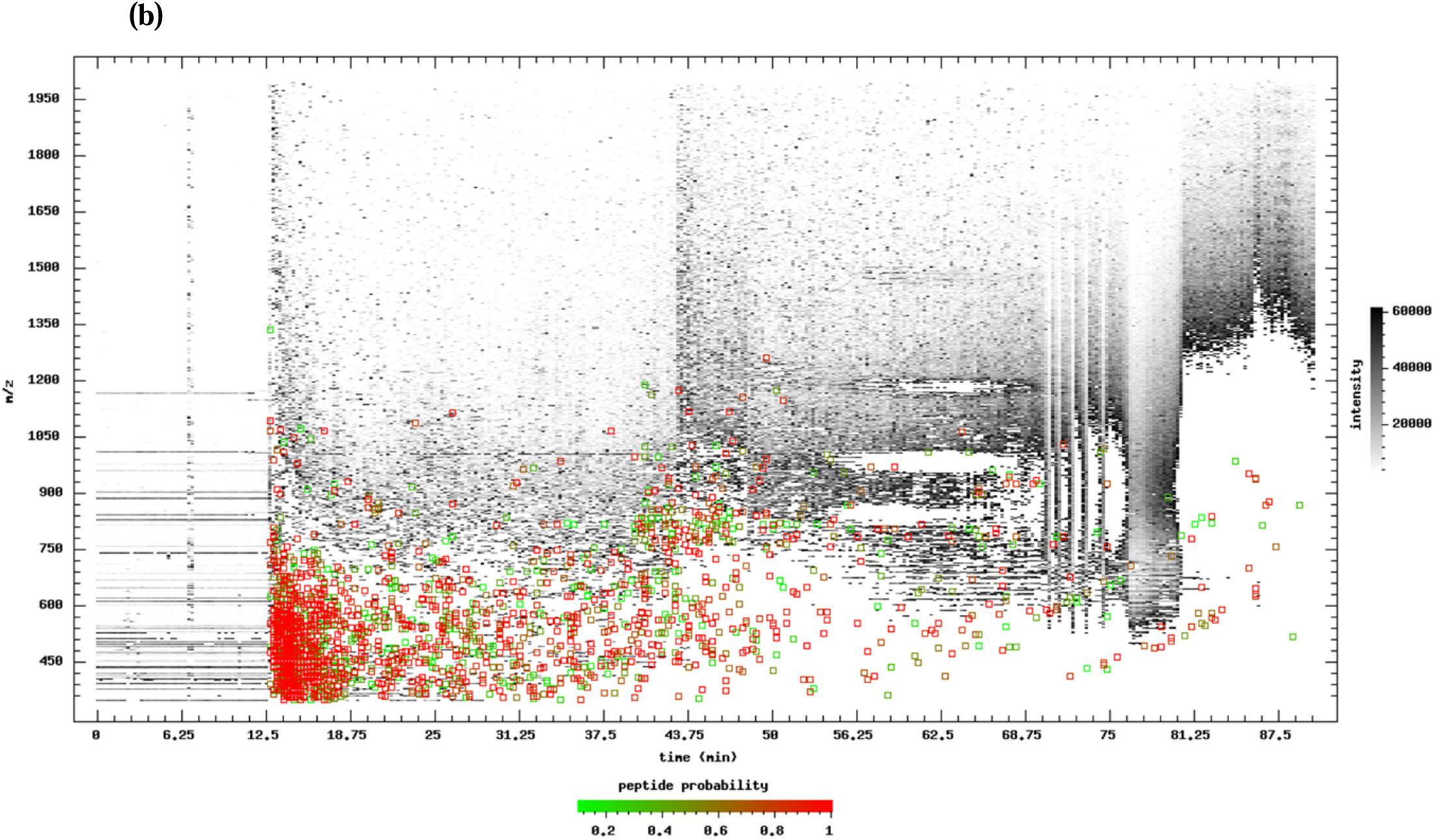
(a) Nano-HPLC chromatogram and (b) total ion spectra of the whey protein peptides.

**Figure 3.2 a.**
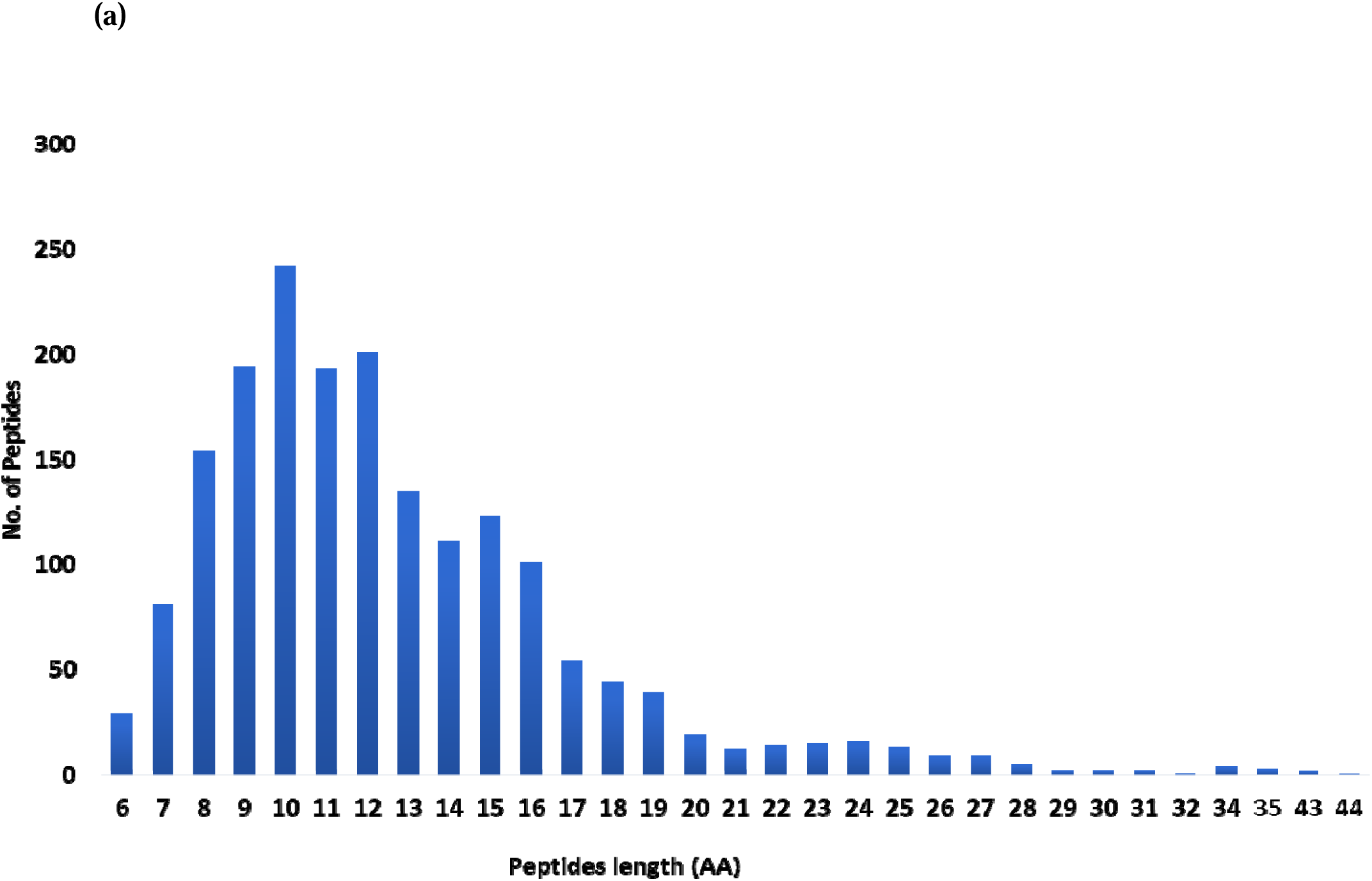
Numbers of peptides amino acid residues.

### 3.2. In silico prediction of bioactivity of peptides by Peptide ranker

Identification of peptides within whey protein hydrolysate were a crucial step in understanding its bioactive potential. In our investigation, a comprehensive analysis revealed the successful identification of a total of 2,883 peptides in library. Among these mostly peptide was observed to be bioactive in nature according to bioactivity score, but only we selected 40 peptides classified as bioactive peptides, characterized by a probability score ranging from 0.90 to 1, as determined by Peptide Ranker (Fig. 3.2 b, Supplementary-II). Sequence logos were constructed for peptides exhibiting bioactivity scores in the range of 0.90 to 1, indicating cysteine, proline, tryptophan, phenylalanine, methionine, and glycine as the predominant amino acid residues (Fig. 3.2 c).

**Figure 3.2 b.**
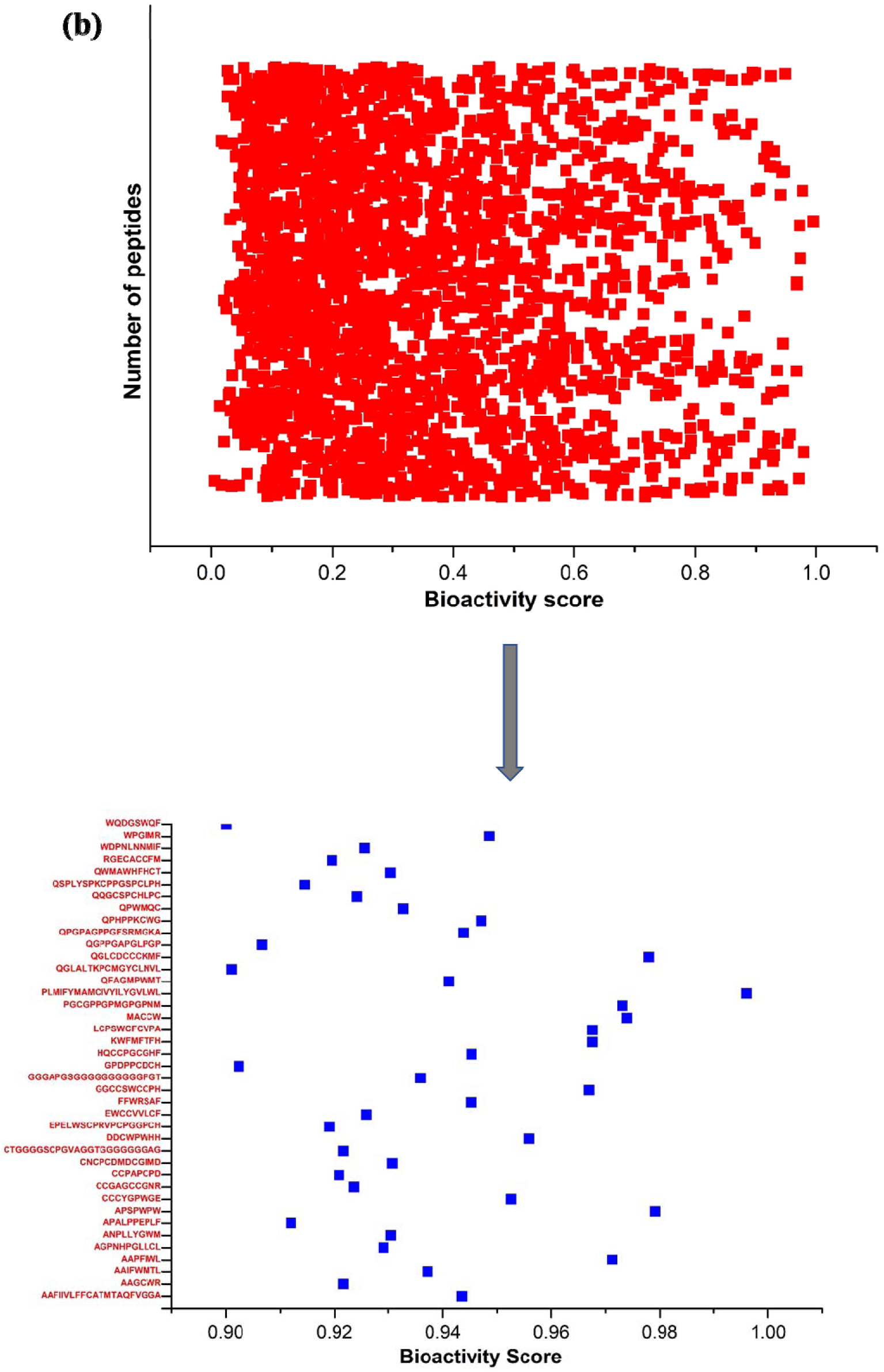
Bioactivity score of identified peptides from whey protein hydrolysate.

**Figure 3.2 c.**
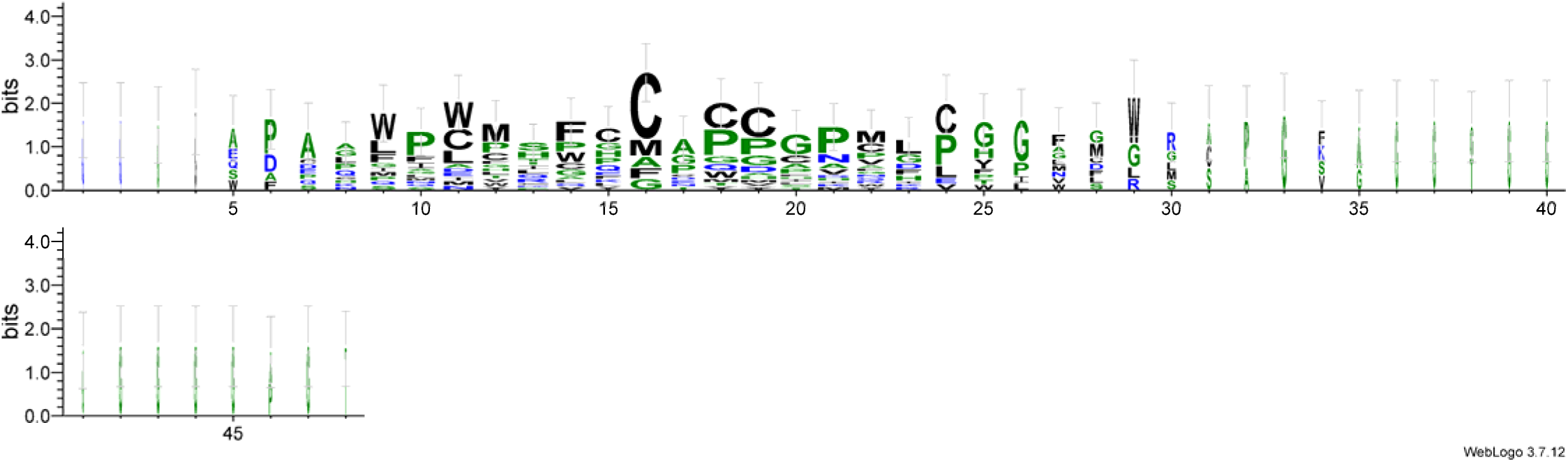
Peptide sequence logos were generated for the sequences predicted with bioactivity scores ranging from 0.90 to 1. The peptide sequence logo for these sequences displayed cysteine, proline, tryptophan, phenylalanine, methionine, and glycine as the dominant residues.

28 peptides predicted as nontoxic out of 40 by Toxinpred (Fig. 3.2 d, Supplementary-II). This classification underscores the potential bioactivity of these peptides, suggesting their significance in various physiological processes or functional applications. One study proposed the *in-silico* prediction of novel peptides, potentially bioactive and bio-accessible from chicken hydrolysate (Xiao et al., 2022), while another study conducted by Coscueta, (2022) utilized a peptide ranker to predict bioactive peptides from casein hydrolysate.

**Figure 3.2 d.**
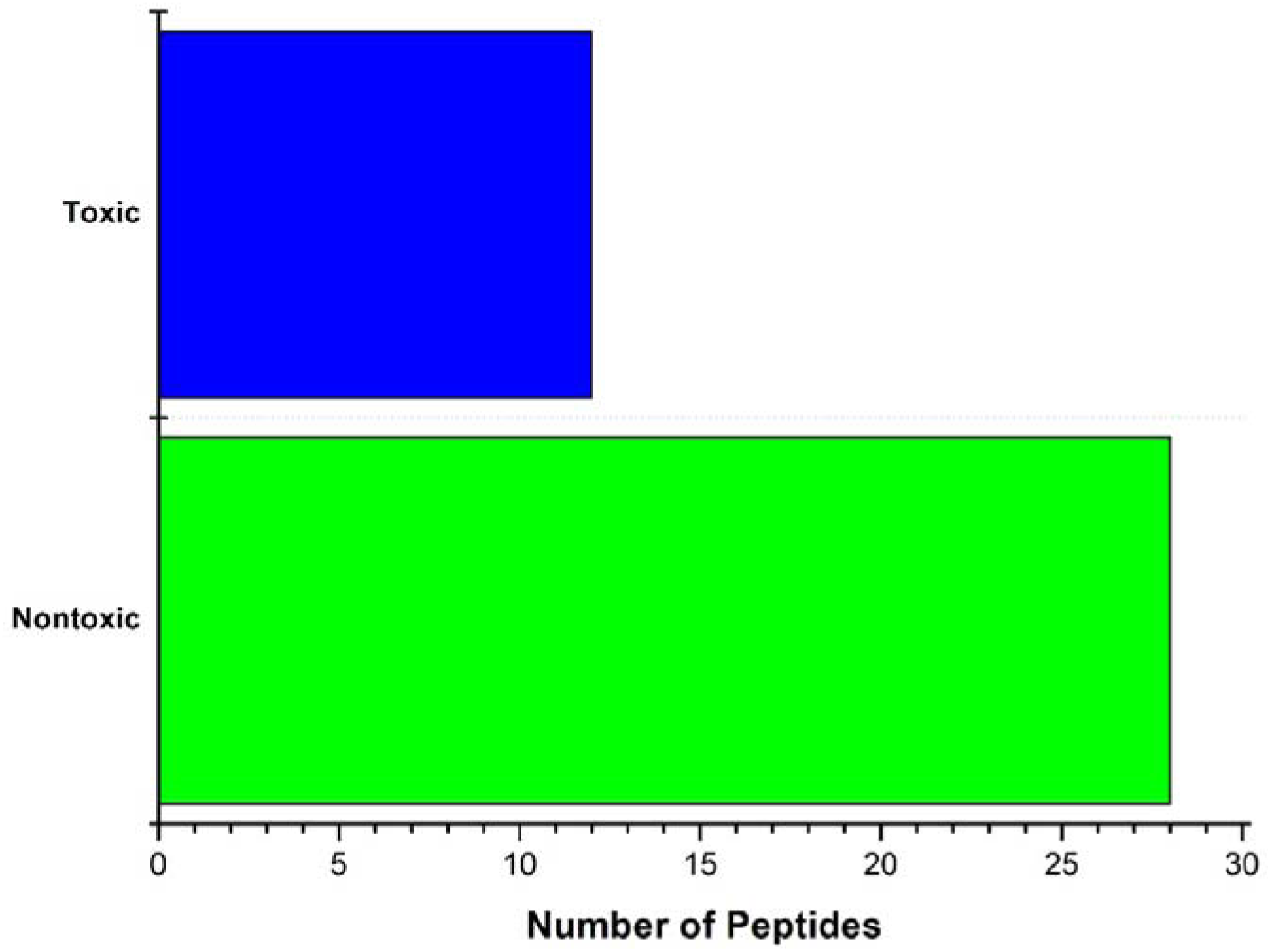
Toxic and nontoxic peptide predicted with bioactivity scores ranging from 0.90 to 1.

The identification of these bioactive peptides within whey protein hydrolysate holds significant implications for its potential health benefits.

### 3.3. Prediction of total molecular weight of bioactive peptides

*In silico* prediction of molecular weight (MW) for peptides plays a crucial role in various fields such as bioinformatics, pharmaceuticals, and proteomics. Molecular weight is a key factor in determining the bioavailability of peptides. Peptides with low molecular weights are more likely to be absorbed and distributed effectively within the body (Romay et al., 2003). In our study, MW predictions of peptides offer that all peptides fell within the range of 500 to 2550 Da (Fig. 3.3). This range is significant as it encompasses peptides of varying sizes, from small bioactive peptides to larger protein fragments. Peptides within this MW range are of particular interest due to their potential biological activities and therapeutic applications.

**Figure 3.3.**
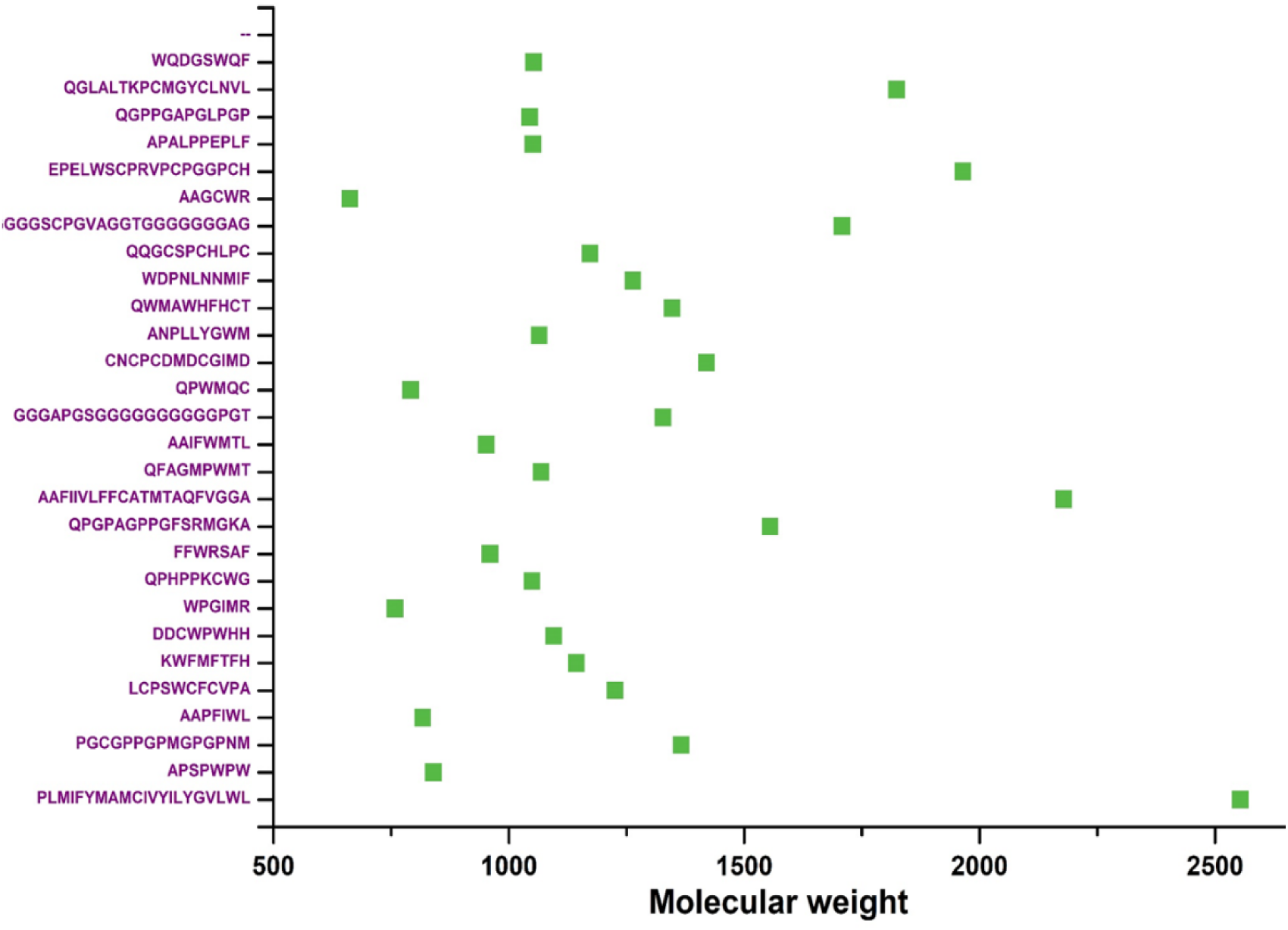
Molecular weight of identified peptides.

### 3.4. Molecular docking analysis of the pancreatic lipase and alpha amylase with selected bioactive peptides

Molecular docking analyses using AutoDock Vina provided detailed insights into the interactions of pancreatic lipase (PL) and alpha-amylase with bioactive peptides derived from WP-PTC, highlighting their potential therapeutic applications. Pancreatic lipase, essential for hydrolyzing dietary fats, features an active site in the N-terminal domain comprising a catalytic triad (S153, D177, H264) and key substrate-binding residues (F78, I79, H152, F216, W253, R257). This site is regulated by a flexible flap that governs enzyme activation by adopting open or closed conformations (Hermoso et al., 1996).

Among the 28 bioactive peptides evaluated, binding energies ranged from 121.9 to −6.5 kcal/mol, with WPGIMR and WQDGSWQF demonstrating the strongest binding energies of −6.5 and −6.1 kcal/mol, respectively (Fig. 3.4. a). WPGIMR interacted with PL via six hydrogen bonds and three pi bonds, while WQDGSWQF showed even more robust interactions with eight hydrogen bonds and eight pi bonds (Fig. 3.4. b & c). These interactions indicate strong structural stability within the enzyme’s binding pocket, suggesting that these peptides could serve as effective inhibitors or modulators of pancreatic lipase activity.

**Figure 3.4.**
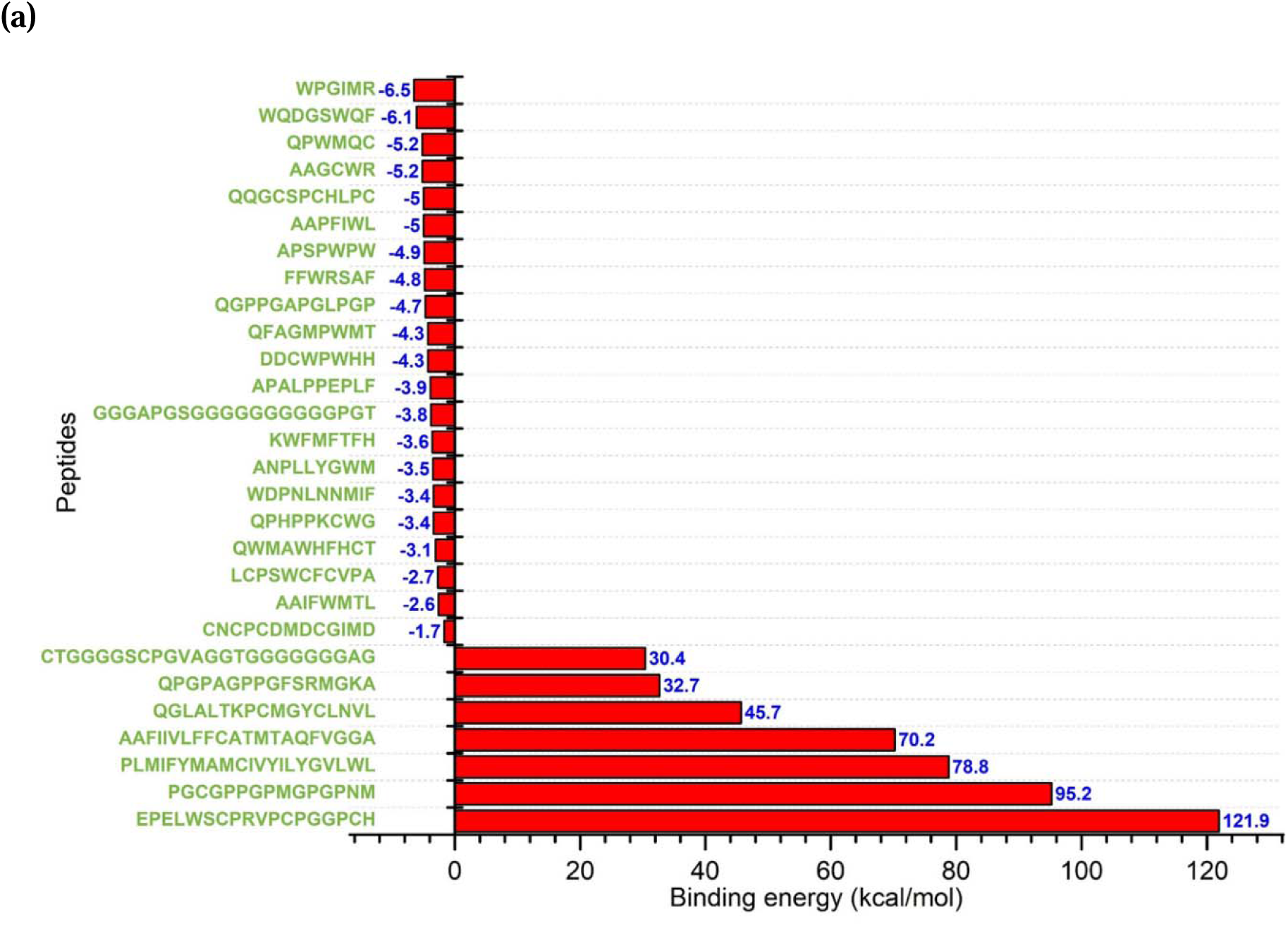
(a) Binding energy of the interacted bioactive peptides with pancreatic lipase.

**Figure 3.4. b.**
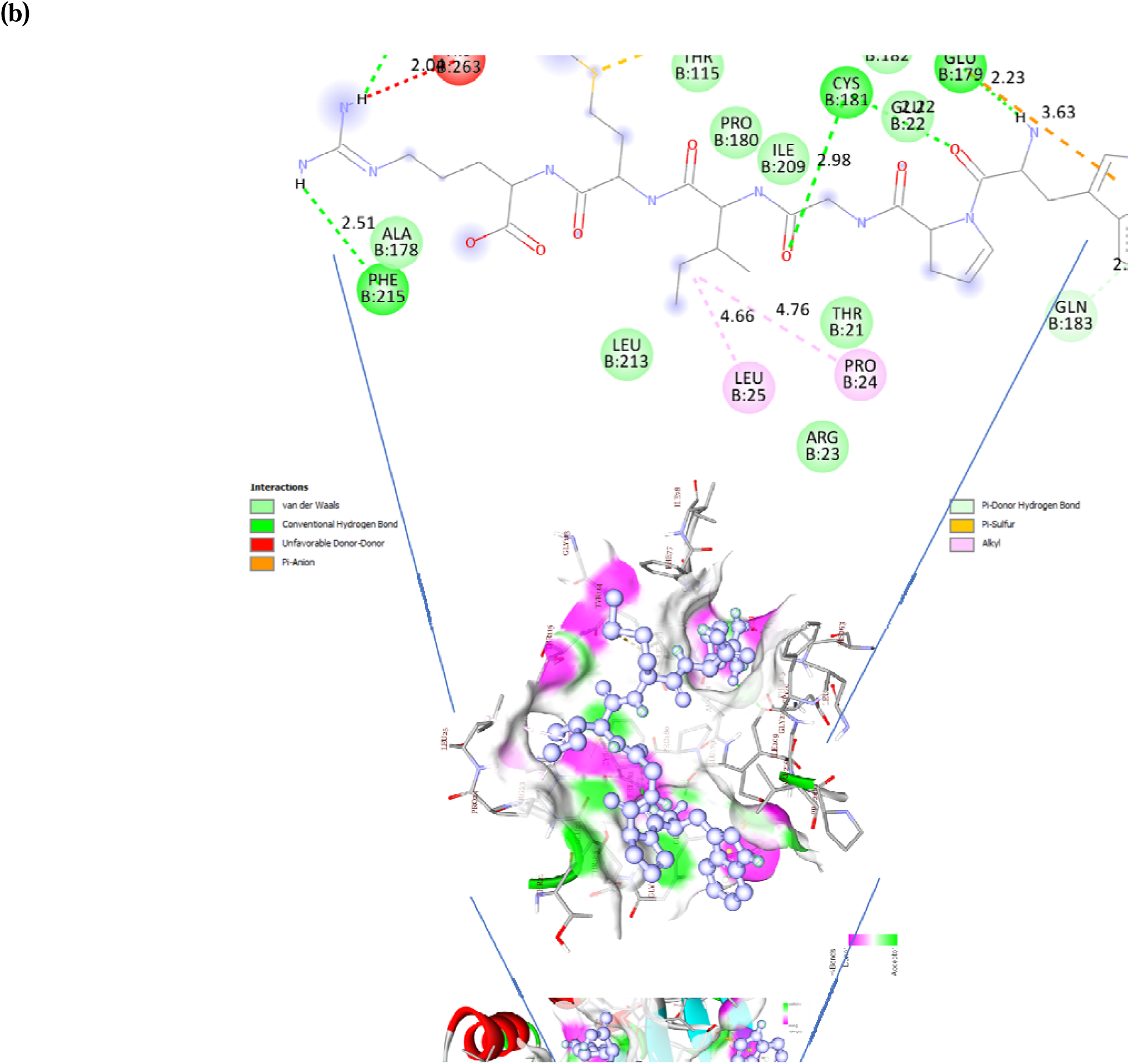
2-D binding analysis of the bioactive peptide WPGIMR with pancreatic lipase.

**Figure 3.4. c.**
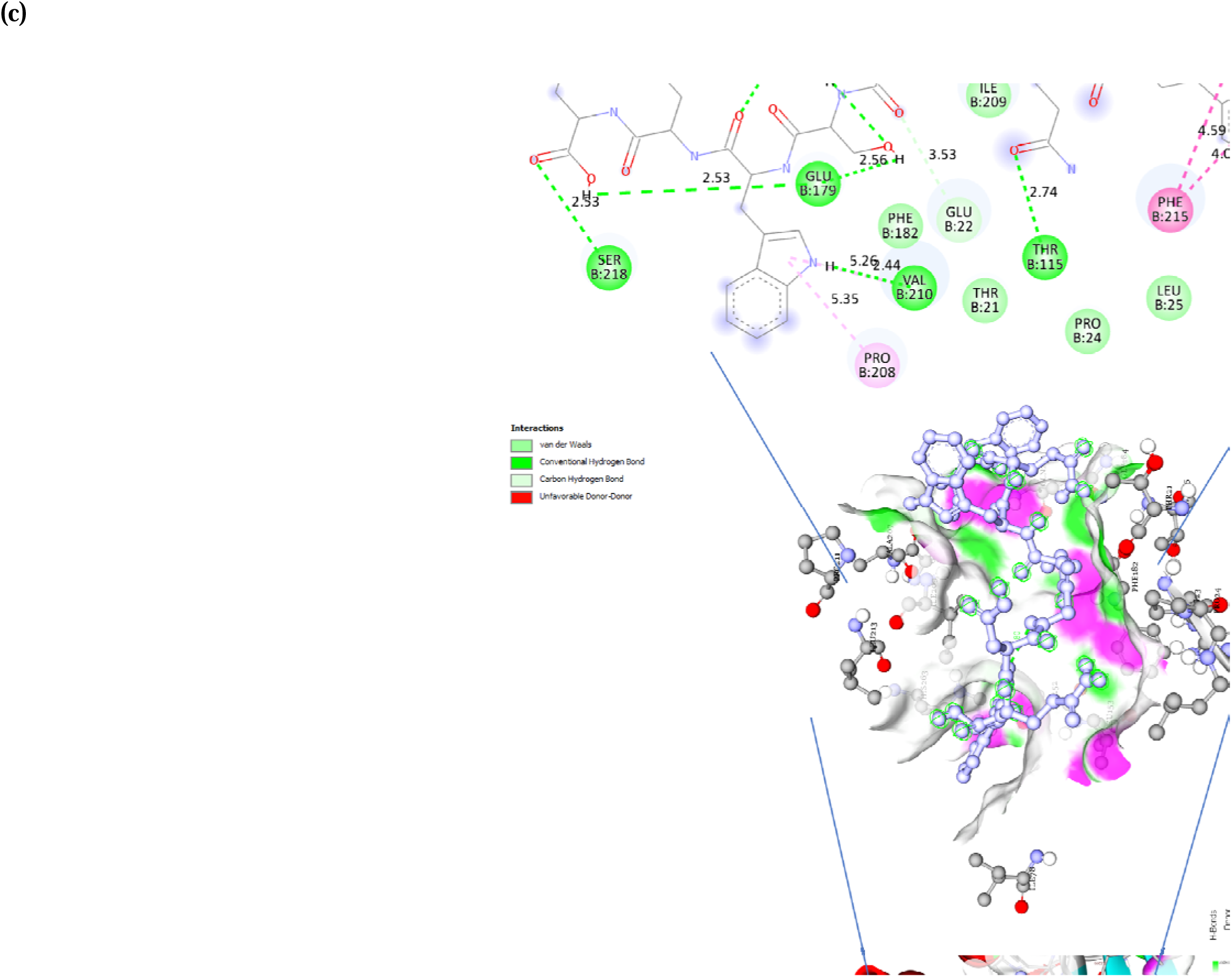
2-D Binding analysis of the bioactive peptide WQDGSWQF, with pancreatic lipase.

Hirose, (2024) identified two anti-obesity peptides, LDQW and LKPTPEGDLEIL, from whey protein using LC-MS, which showed promise in combating obesity and metabolic disorders. Similarly, camel whey protein-derived peptides such as PAGNFLPPVAAAPVM, MLPLMLPFTMGY, and LRFPL demonstrated inhibitory potential against PL through molecular docking analysis (Baba et al., 2021). These peptides interacted with PL primarily via hydrogen bonds, salt bridges, and hydrophobic interactions, highlighting their potential to destabilize the open-lid conformation of the enzyme.

Validated peptides like VPPR, LADR, LSPR, and TVGPR further exhibited uncompetitive inhibition of PL, emphasizing the role of structural motifs and interaction types for enzyme regulation. Other studies identified peptides such as MSNFYF (camel casein) and MMML (cow casein) that bind at the active site of PL, confirming their regulatory activity (Mudgil et al., 2022). Additionally, plant-derived peptides from sources like cocoa beans, yellow field peas, and pinto beans revealed the importance of non-polar residues in forming stable complexes with PL (Mudgil et al., 2018; Coronado-Cáceres et al., 2020; Awosika & Aluko, 2019; Ngoh et al., 2017). These findings collectively demonstrate the potential of bioactive peptides as natural pancreatic lipase inhibitors, offering promising strategies for managing obesity and associated metabolic disorders.

Molecular docking analyses of α-amylase revealed significant interactions between bioactive peptides and the enzyme’s key active sites. α-Amylase consists of three subunits—A, B, and C—with the primary active sites Asp197, Glu233, and Asp300 located in subunit A, along with calcium-binding (Asp100, Arg158, Asp167, His201) and chloride-binding (Arg195, Asn298, Arg337) domains, which are critical for enzymatic activity (Esfandi et al., 2022).

In our study, binding energies of peptides with α-amylase ranged from 71 to −8.4 kcal/mol, indicating diverse affinities (Fig. 3.4. d). Notably, peptides AAPFIWL and WQDGSWQF exhibited the strongest binding energies of −8.4 and −8.1 kcal/mol, respectively (Fig. 3.4. d). AAPFIWL formed 5 hydrogen bonds and 7 pi bonds, contributing to stability within the enzyme’s binding site. Similarly, WQDGSWQF established 8 hydrogen bonds and 8 pi bonds, highlighting its robust interactions with α-amylase (Fig. 3.4. e & f). These diverse modes of interaction demonstrate the peptides’ potential as α-amylase inhibitors.

**Figure 3.4. d.**
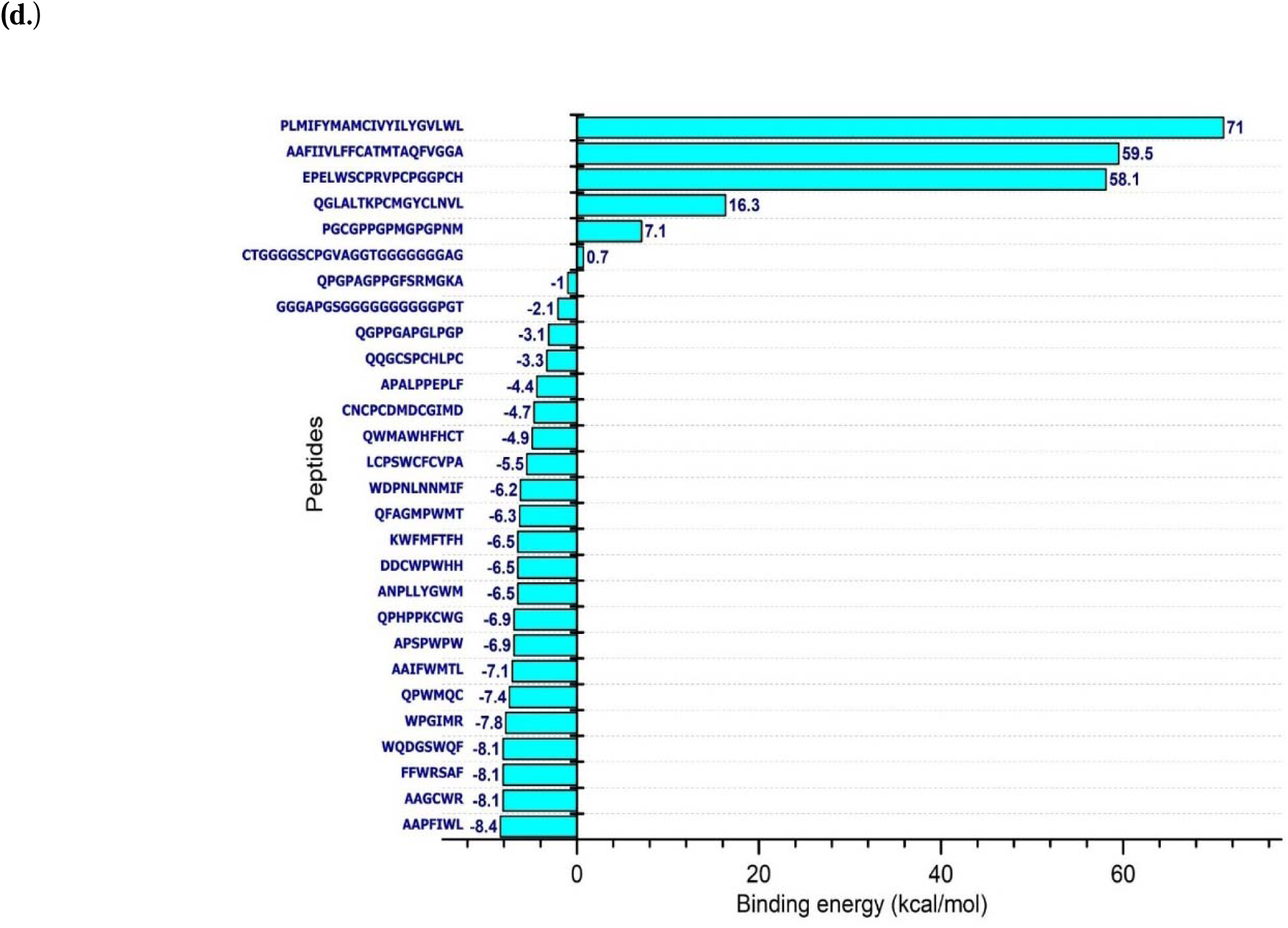
Binding energy of the interacted bioactive peptides with α-amylase.

**Figure 3.4. e.**
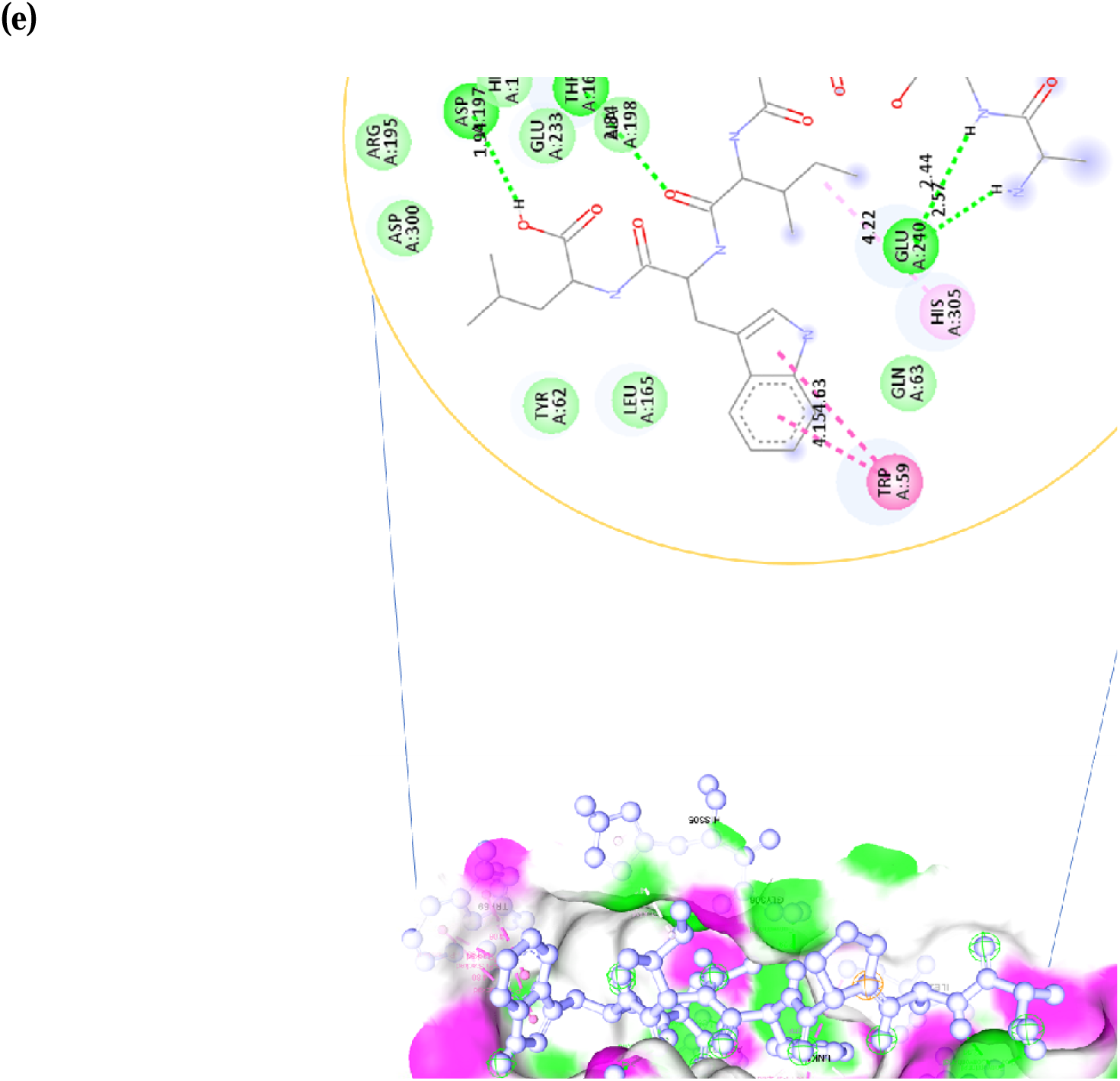
2-D Binding analysis of the bioactive peptide AAPFIWL with Alpha amylase.

**Figure 3.4. f.**
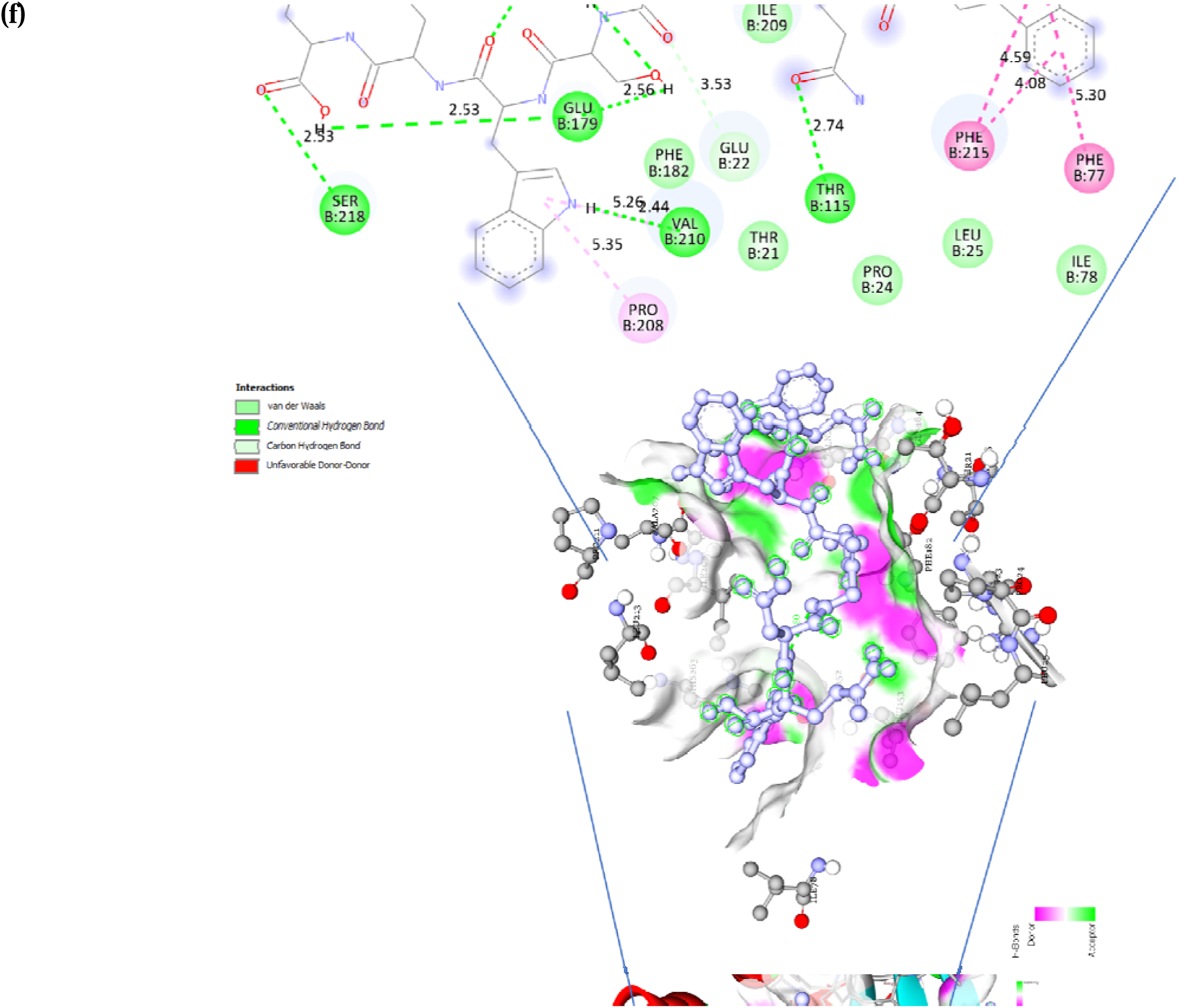
2-D binding analysis of the bioactive peptide WQDGSWQF, with Alpha amylase.

On of the study, Mudgil, (2024) identified peptides IMEQQQTEDEQQDK from camel milk and DQHQKAMKPWTQPK from goat milk, which showed strong binding affinity towards α-amylase. Another study by Su, (2024) identified QEPVPDPVRGL from camel casein protein, which exhibited potent antidiabetic activity by effectively occupying the enzyme’s active cavities through hydrogen bonds, salt bridges, and pi-alkyl interactions. Furthermore, Singh, (2024) demonstrated the enzymatic hydrolysis of whey protein from the Himalayan goat breed “Gaddi” using alcalase, flavourzyme, and their combination. This process yielded bioactive peptides TPEVDKEALEK (−5.7 kcal/mol), DDSPDLPK (−6.5 kcal/mol), and EMPFPK (−7.0 kcal/mol). These peptides exhibited strong and efficient interactions with α-amylase, further highlighting their potential as bioactive molecules for therapeutic applications.

Collectively, these results emphasize the critical role of bioactive peptides in modulating α-amylase activity, underscoring their potential in developing functional foods and therapeutic agents for managing metabolic disorders.

In summary, these findings underscore the potential of select bioactive peptides sourced from whey protein hydrolysates to interact with and potentially influence the activities of pancreatic lipase and alpha-amylase. Insight into the molecular mechanisms governing these interactions is essential for the identification of innovative therapeutic agents or functional food constituents targeting lipid and starch metabolism.

### 3.5. Peptide sequence logos for pancreatic lipase and α-amylase binding peptide

The predicted binding energies of peptides interacting with the active sites of pancreatic lipase and α-amylase ranged from −3.1 to −8.4. Sequence logo analysis of peptides binding to pancreatic lipase indicated tryptophan, methionine, phenylalanine and proline as the most dominant hydrophobic and aromatic residues. In contrast, sequence logo analysis of peptides binding to α-amylase identified tryptophan, leucine and proline as the predominant aromatic and hydrophobic residues (Fig. 3.5a & b).

**Figure 3.5.**
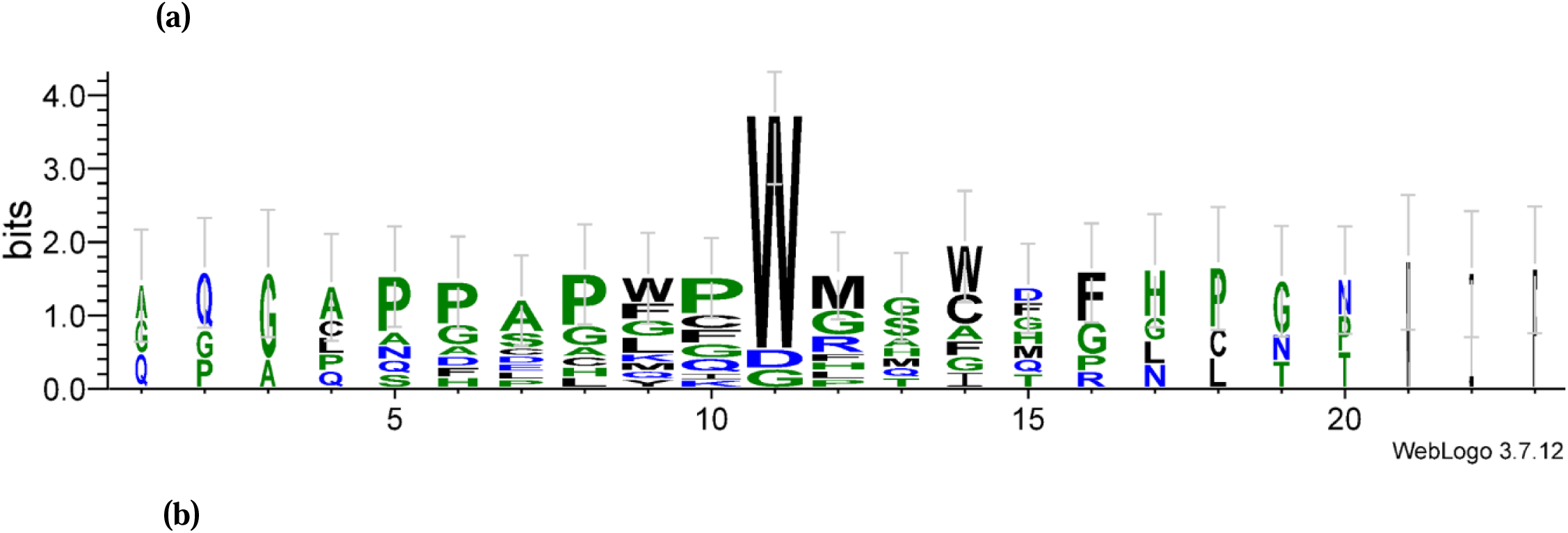
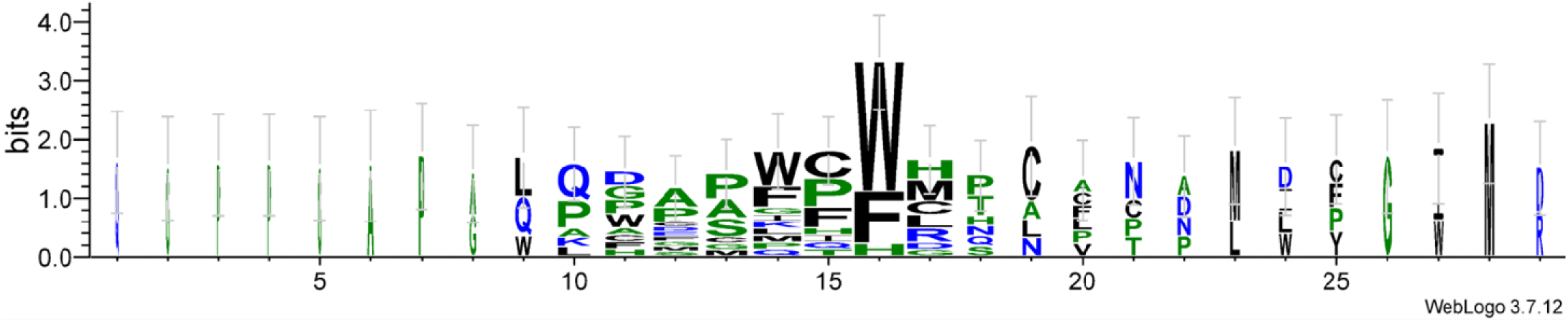
The binding energy, ranging from −3.1 to −8.4, of peptide sequence logos was generated for the sequences predicted to bind with the pancreatic lipase and α-amylase pockets. (a) The peptide sequence logo for the pancreatic lipase binding peptide sequences displayed tryptophan, methionine, phenylalanine and proline as the dominant residues. (b) The peptide sequence logo for the predicted α-amylase binding peptide sequences highlighted tryptophan, leucine and proline as the predominant the dominant residues.

## Conclusion

This study successfully identified 2,883 peptides in whey protein hydrolysates using HR-LC/MS guided library preparation, with 28 peptides predicted as bioactive based on a high probability score (0.9–1) from PeptideRanker. These peptides, predominantly 8 to 13 amino acids in length, demonstrated significant inhibitory potential against pancreatic lipase and alpha-amylase through molecular docking analysis. For pancreatic lipase, binding energies ranged from 121.9 to −6.5 kcal/mol, with WPGIMR and WQDGSWQF showing the strongest affinities, forming multiple hydrogen and pi bonds that underscored their robust interactions with the enzyme. Similarly, alpha-amylase docking revealed binding energies between −8.4 and −7.1 kcal/mol, with AAPFIWL and WQDGSWQF exhibiting the highest affinities, demonstrating diverse interaction modes and structural stability. The findings highlight the potential of whey protein-derived peptides as inhibitors of key metabolic enzymes involved in lipid and carbohydrate digestion. These results contribute valuable insights into the molecular mechanisms of enzyme inhibition, paving the way for their application as functional food components or therapeutic agents for managing metabolic disorders such as obesity and diabetes. Future studies should address limitations by exploring toxicity, pharmacokinetics, and conducting *in vitro* and *in-vivo* trials to validate the safety and efficacy of these bioactive peptides, enabling their practical application in dietary and pharmaceutical interventions.

## AUTHOR CONTRIBUTIONS

Manish Singh Sansi: Conceptualization, Data curation, Methodology, Visualization, writing– original draft | Daraksha Iram: Data curation, Visualization, writing – review | Sudarshan Kumar: Writing – review and editing | Suman Kapila: Provide instruments, writing – review and editing | Kamal Gandhi: Provide instruments, writing – review and editing | Sunita Meena: Conceptualization, Project administration, writing – review and editing

## ACKNOWLEDGMENTS

We are grateful to the National Dairy Research Institute, India, for providing the necessary facilities to carry out this research. We also thank the Indian Council of Medical Research (ICMR) for awarding the research fellowship during this study period. Additionally, we acknowledge the Sophisticated Analytical Instrument Facility (SAIF), IIT Bombay, for providing access to the HR-LC/MS facility for peptide sequencing.

## Data availability statement

The data and materials supporting our findings are available from the corresponding author upon reasonable request.

